## Supplementary Information for "Dynamic basis of lipopolysaccharide export by LptB_2_FGC"

**Supplementary Figures**

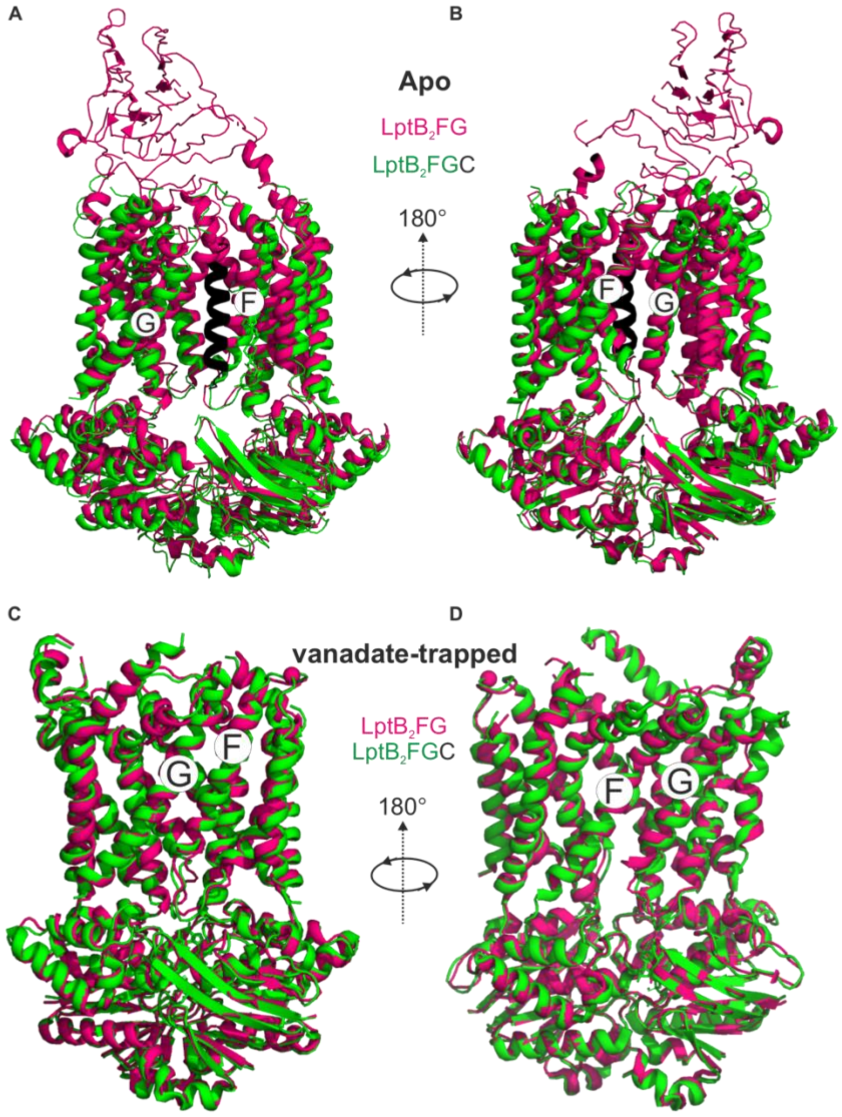


**Figure S1. $LptB_2FG$ and $LptB_2FGC$ structures in the apo and vanadate-trapped states.** (A) *E. coli* $LptB_2FG$ (magenta, PDB ID: 6MHU) and $LptB_2FGC$ ($LptB_2FG$ subunits in green and LptC in black, PDB ID: 6MI7) structures in the apo state are overlaid. The gating helices named as G and F moves apart upon binding LptC. A view from the other lateral gate is shown in (B). (C) *E. coli* $LptB_2FG$ (magenta, PDB ID: 6MHZ) and $LptB_2FGC$ (PDB ID: 6MI8) structures in the vanadate-trapped states are overlaid. The TM-LptC is not resolved in $LptB_2FGC$ and the structures are nearly identical with an overall RMDS of 0.99 Å.

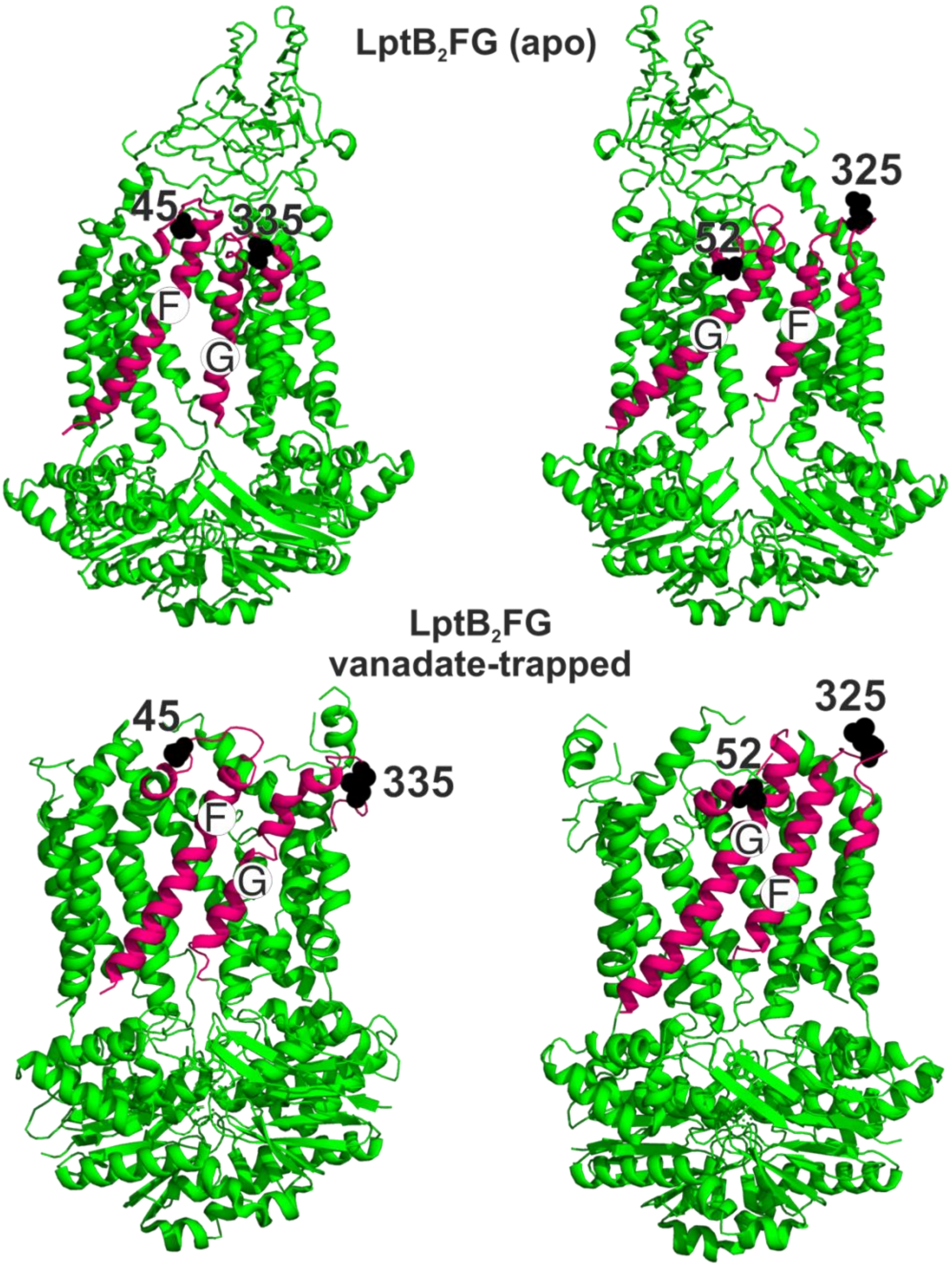


**Figure S2. Spin labeled positions in $LptB_2FG$.** The two lateral gates (magenta) and the labeled positions (black spheres) are highlighted on the apo (PDB ID: 6MHU) and vanadate-trapped (PDB ID: 6MHZ) $LptB_2FG$ structures (top and bottom, respectively).

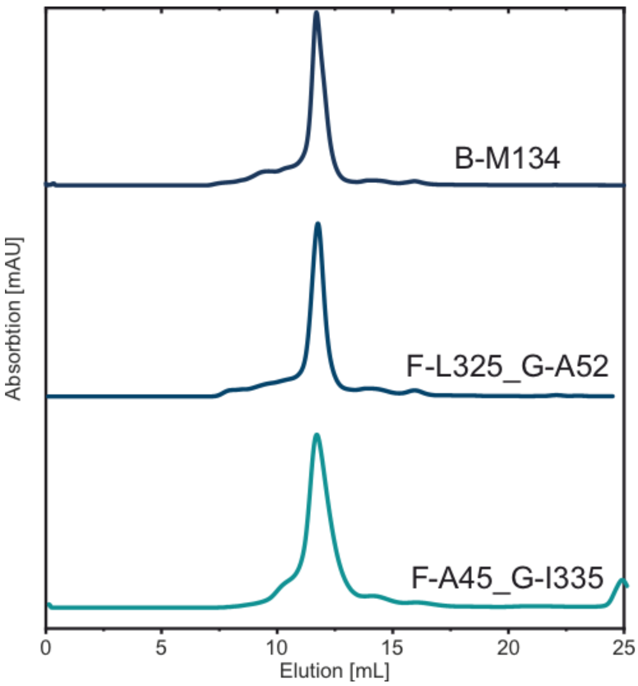


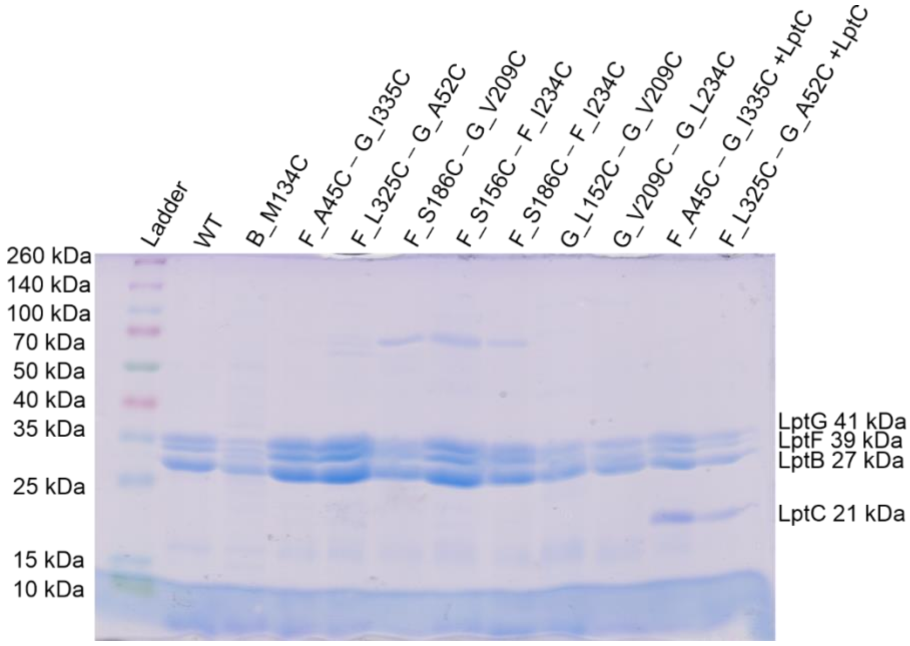


**Figure S3. Size-exclusion chromatography (SEC) and SDS-PAGE for WT and spin labeled cysteine variants of LptB$_2$FG and LptB$_2$FGC**. (Top panel) Typical SEC profiles for selected spin labeled variants as indicated are shown. (Bottom panel) SDS-PAGE gel. Proteins were collected from SEC column and the molecular weight of the subunits are indicated (LptG 41 kDa, LptF 39 kDa, LptB 27 kDa and LptC 21 kDa).

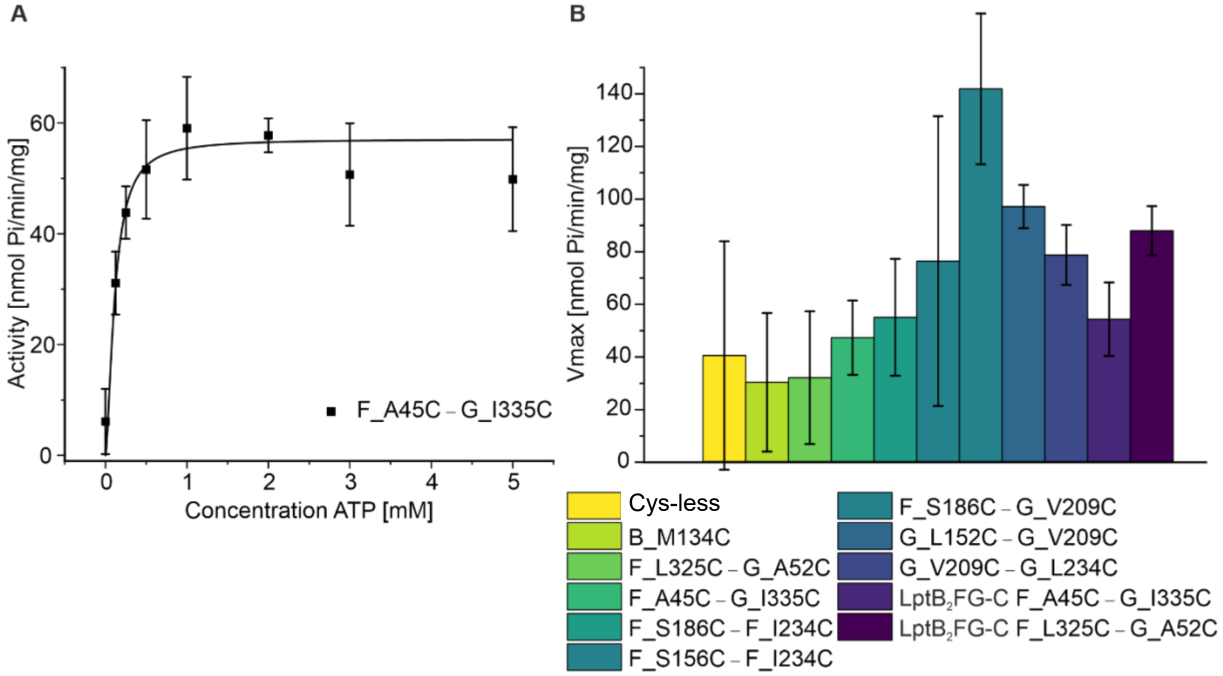


**Figure S4. ATPase assay for spin labeled variants of $LptB_2Fg$ and $LptB_2FGC$**. The functionality of the spin labelled proteins were characterized using ATPase assay and typical reaction curve is shown (left). The corresponding $V_{max}$ values are shown (right). All the variants actively hydrolysed ATP, in line with the conformational changes observed from PELDOR/DEER experiments following vanadate-trapping (Figures 2-6). As the variants are spin labelled at different positions, a quantitative comparison between the values is omitted.

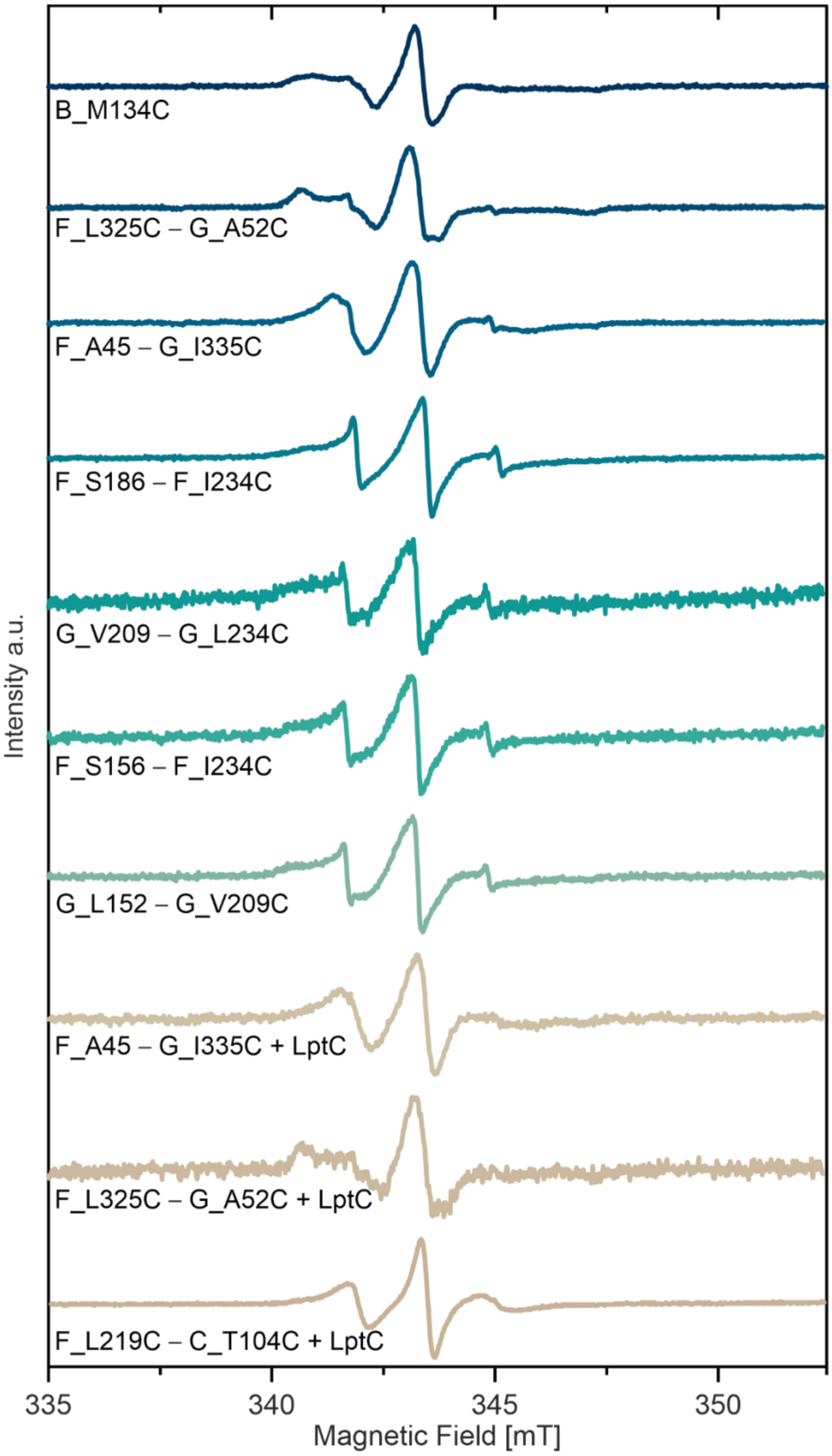


**Figure S5. Room temperature continuous wave ESR spectroscopy of MTSL labeled variants in micelles**. Spectra for spin labelled LptB$_2$FG and LptB$_2$FG-C variants in DDM micelles are shown.

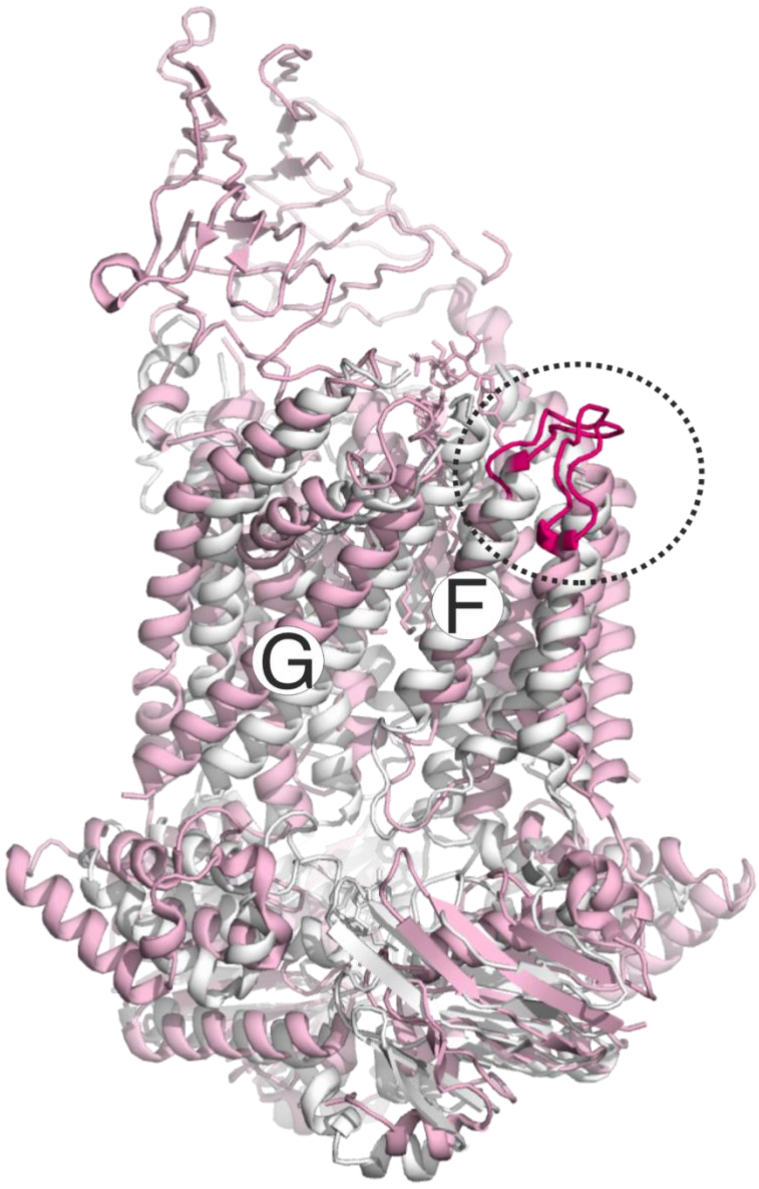


**Figure S6. Conformation of the loop carrying the spin labeled position 325 in LptF at the second lateral gate.** $LptB_2FG$ apo (pink, PDB ID: 6MHU) and vanadate-trapped (grey, PDB ID: 6MHZ) structures are overlaid. The loop carrying the spin labelled position L325 is highlighted (magenta), which has a rather similar conformation in the two structures.

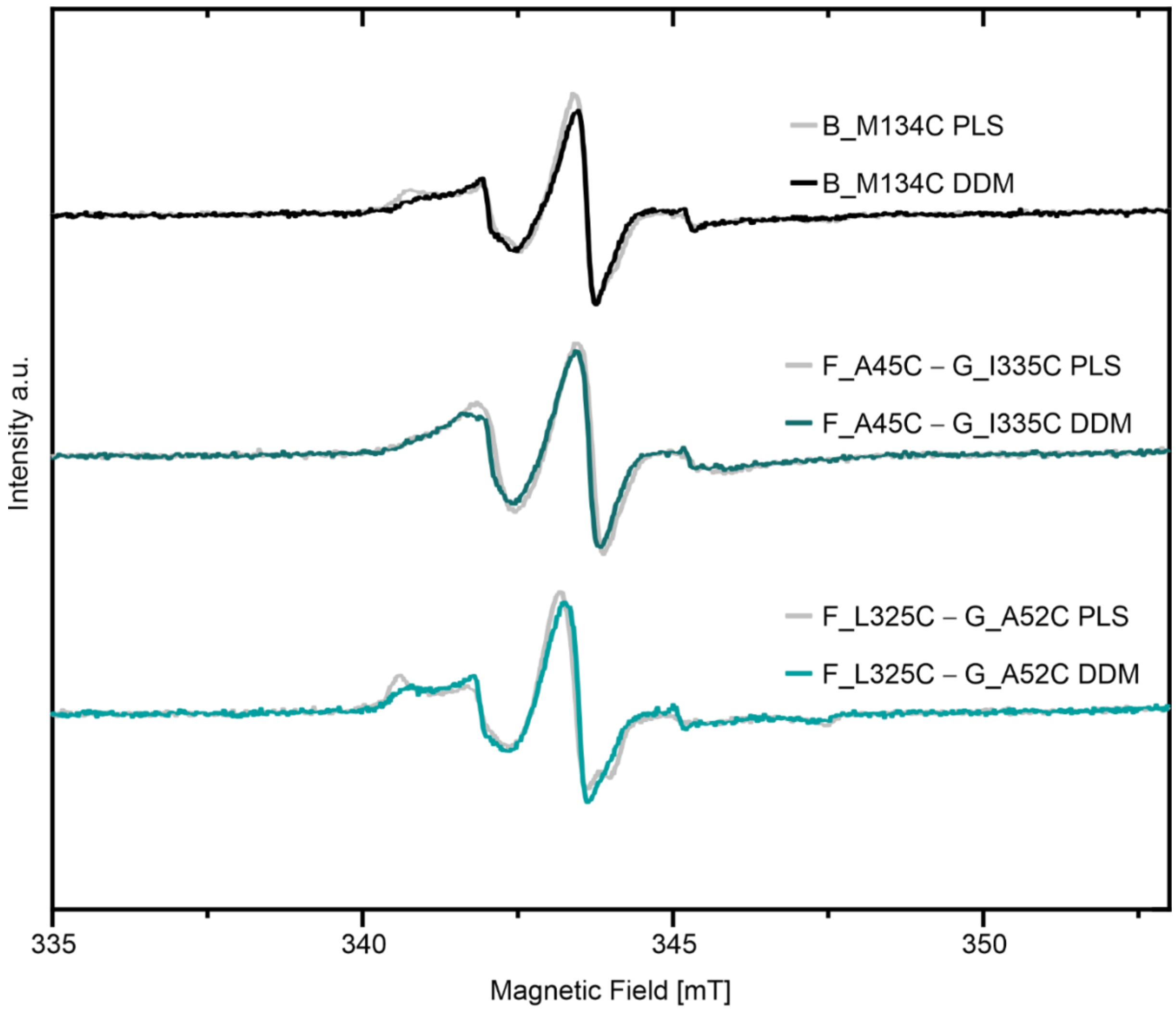


**Figure S7. Comparison of room temperature continuous wave ESR spectroscopy of MTSL labeled variants between DDM micelles and proteoliposomes (PLS)**. Corresponding spectra for spin labelled $LptB_2FG$ variants in liposomes and DDM micelles are shown.

| Variant | Protein [µM] (±10%) | Spin [µM] (±10%) | Labelling efficiency [%] (±15%) |
|---|---|---|---|
| B_M134C PLS | 60 | 125 | 96 |
| F_A45C – G_I335C | 37 | 72 | 98 |
| F_L325C – G_A52C | 37 | 69 | 93 |
| F_S186C – G_V209C | 31 | 70 | 112 |
| F_S156C – F_I234C | 70 | 130 | 93 |
| F_S186C – F_I234C | 31 | 61 | 99 |
| G_L152C – G_V209C | 46 | 87 | 95 |
| G_V209C – G_L234C | 34 | 64 | 94 |
| F_A45C – G_I335C + LptC | 29 | 58 | 100 |
| F_L325C – G_A52C + LptC | 31 | 62 | 100 |

**Table S1. Spin labelling efficiency for the investigated cysteine variants.** A 10% error is estimated for the protein and spin concentrations.
